# Prediction of Outcome from Spatial Protein Profiling of Triple-Negative Breast Cancers

**DOI:** 10.1101/2025.04.18.649541

**Authors:** Ali Foroughi pour, Te-Chia Wu, Jan Martinek, Santhosh Sivajothi, Paul Robson, Karolina Palucka, Jeffrey H. Chuang

## Abstract

In tumors, reciprocal spatial interactions between immune cells, their mediators, the extracellular matrix, and mutated neoplastic cells impact all aspects of treatment resistance. The operational mechanisms of these interactions are foundational for developing insights and targets for cancer therapy and prevention. Spatial quantification of the tumor microenvironment system from image data has untapped potential for patient stratification. Here, we present SparTile, a powerful computational approach for analysis of multiplex proteomics images to reveal clinically relevant structural organization in the tumor microenvironment. SparTile enables the robust unbiased identification and characterization of tumor microenvironments based on spatial relationships among protein markers without the need for cell segmentation or classification. Applied to tissues of patients with triple negative breast cancer (TNBC), an aggressive subtype of breast cancer, SparTile identified repeatable microenvironments with specific cellular relationships. These included microenvironments characterized by risk markers such as Ki67+ (p-value= 0.052) and vimentin+ (p-value<0.01) tumor cells, which correlate with poor survival. Furthermore, myeloid markers in an MX1 positive tumor environment correlate with shorter survival (p-value=0.04). Finally, we identified a relative distance between tumor and myeloid cell markers as a strong prognostic risk factor in multivariate Cox models (p-value=0.01). This distance metric was externally validated on two datasets of breast cancer multiplex images (p-values<0.01). Thus, unbiased protein-based and segmentation-free spatial analysis is effective for identifying clinically relevant biomarkers from multiplex tumor images and identifies novel predictive biology.

## Introduction

The tumor microenvironment is a complex amalgamation of diverse interacting components, such as individual cells, structural components, and signaling proteins. These components impact pro- and anti-tumor immune activity, progression, metastasis, and survival risk. While recent studies have shown the importance of tumor-immune interactions in response to therapy and risk stratification, common descriptors of immune activity such as the abundances of tumor-infiltrating lymphocytes (TILs) and tumor associated macrophages, explain only a limited subset of the behaviors that pervade the tumor microenvironment (Bianchini et al, 2022, Zagami and Carey 2022).

Multiplex antibody-based imaging techniques, such as imaging mass cytometry (IMC) now enable simultaneous measurement of dozens of epitopes at subcellular resolution, but computational approaches for interpretation of these data types are lacking. Most current approaches (e.g., Ali et al, 2020, Jackson et al, 2020, Shiao et al, 2024) are predicated on cell segmentation and classification, but these steps are challenging and have fundamental limitations in 2D. Furthermore, categorizing cells into a limited number of classes obfuscates the heterogeneity of differentiated and regulated states within each cell class. To address these challenges, we have developed SparTile, a novel computational approach for analyzing multiplex protein images. Inspired by deep learning approaches in digital pathology, SparTile breaks protein images into smaller tiles and uses sparse non-negative matrix factorization to generate low dimensional tile representations. These tile representations are interpretable and can be used to robustly identify microenvironments (MEs) within and across tumors.

To demonstrate the power of SparTile, we have studied tissue samples from patients with triple negative breast cancer (TNBC), which account for 15-20% of all breast cancers and carry the worst prognosis across all breast cancers. TNBCs are heterogeneous and an improved understanding of the tumor-immune ecosystem is necessary to develop prognostic markers of risk and response to therapy (Bianchini et al, 2022). Using SparTile, we identify and confirm MEs previously shown to correlate with survival, identify and externally validate MEs associated with survival, and distinguish fine spatial arrangements of cells that can be quantitatively associated with patient outcome. Our work demonstrates how segmentation-free analysis of multiplex antibody-based imaging of tumors can be used effectively for clinically-relevant outcome prediction.

## Results

### Multiplex imaging of TNBC tissue microarrays

To investigate tumor-immune interactions in primary TNBC tumors, we generated IMC data from tissue microarray (TMA) spots of 88 tumors. We developed panels enabling imaging of DNA and 45 epitopes. Two (5 um) adjacent slices were registered for each spot to enable profiling of a larger marker set (Fig. 1A). This allowed identification of diverse cellular and microenvironmental markers and thorough characterization of the immune landscape (Figs. 1A,B). The panel for the first slice provided a general characterization of cell composition, while the second panel focused on immune markers. To provide a common feature space for registration of adjacent slices, we generated virtualized hematoxylin and eosin (H&E) images from the IMC data of each slice (methods, Fig. 1C). Registration of adjacent H&E images enabled spatial comparison across all assayed features, though tissue differences between slices constrained cell segmentation-based comparisons (Supplementary Fig. 1). For example, we frequently observed cells within only one of the two slices. We also observed differences in nucleus or cell membrane shapes across slices, not unexpected considering a typical cell size is 10 um and reflecting the fact that slices do not capture whole cells but instead reflect cell cross-sections (Supplementary Fig. 1).

**Figure 1:**
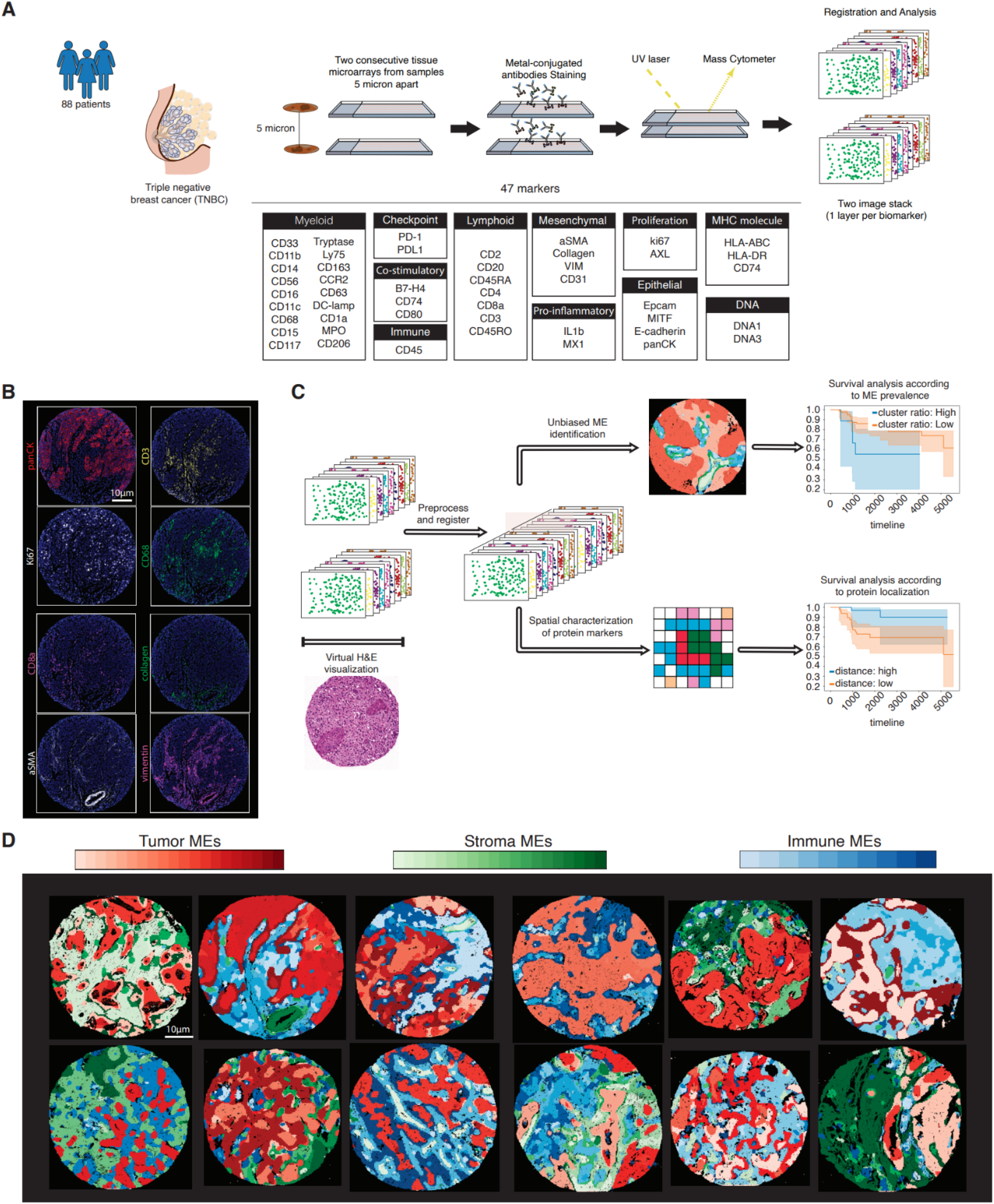
Overview of data acquisition and computational analysis: (A) 5 micron adjacent cuts were stained with metal-conjugated antibodies for multiplex protein imaging using two panels. (B) Multiplex imaging identifies spatial distribution of proteins at subcellular resolutions. (C) Overview of SparTile: Virtualized H&E of IMC images were generated for visual inspection and image registration. Preprocessed images of adjacent cuts were registered by mapping the coordinates of the second image to the first, resulting in 1 multiplex image with 47 markers (2 DNA and 45 proteins). SparTile identifies different MEs via clustering. Fraction of each ME was computed for each patient. Survival analysis was used to identify MEs that are predictive of patient risk. Processed markers of IMC images were used to find their relative spatial distribution. Survival analysis was used to identify spatial relations that are predictive of patient risk. (D) Examples of spatial mappings of SparTile clusters. SparTile identified 20 tumor, 20 stroma, and 10 immune clusters, which downstream analysis interpreted as distinct MEs. These MEs are spatially contiguous, and multiple MEs are present within each spot. Shades of red, green, and blue denote different tumor, stroma, and immune MEs.

### SparTile robustly identifies microenvironments

Given the limitations of segmentation-based analysis, we developed an analysis approach to identify MEs directly from antibody image data. This approach, SparTile (SPARse non-negative matrix factorization for TILE analysis), was designed to robustly identify MEs within and across spatial antibody images. Such MEs can then be studied more deeply for interactions and spatial relationships, as well as for identification of MEs that are predictive of survival (Figs. 1C,D). SparTile first clusters tiles into three general region types: tumor, stroma, and immune (methods, Supplementary Fig. 1). Region type identification occurs at the tile scale (methods), though there may be finer structure within each tile, e.g. low densities of immune cells within predominantly tumor regions (Supplementary Fig. 1). The second step is to distinguish MEs with more specific differences. SparTile uses sparse non-negative matrix factorization (sNMF) to learn low-dimensional representations of tiles within each region type (methods, Fig. 2A). Using these representations, we then learned 20 clusters in the tumor, 20 clusters in the stroma, and 10 clusters in the immune region type (methods, Figs. 1D, 2B, and 2C). Different markers characterize each cluster, indicating that SparTile effectively distinguishes different MEs (methods, Figs. 1D and 2B).

**Figure 2:**
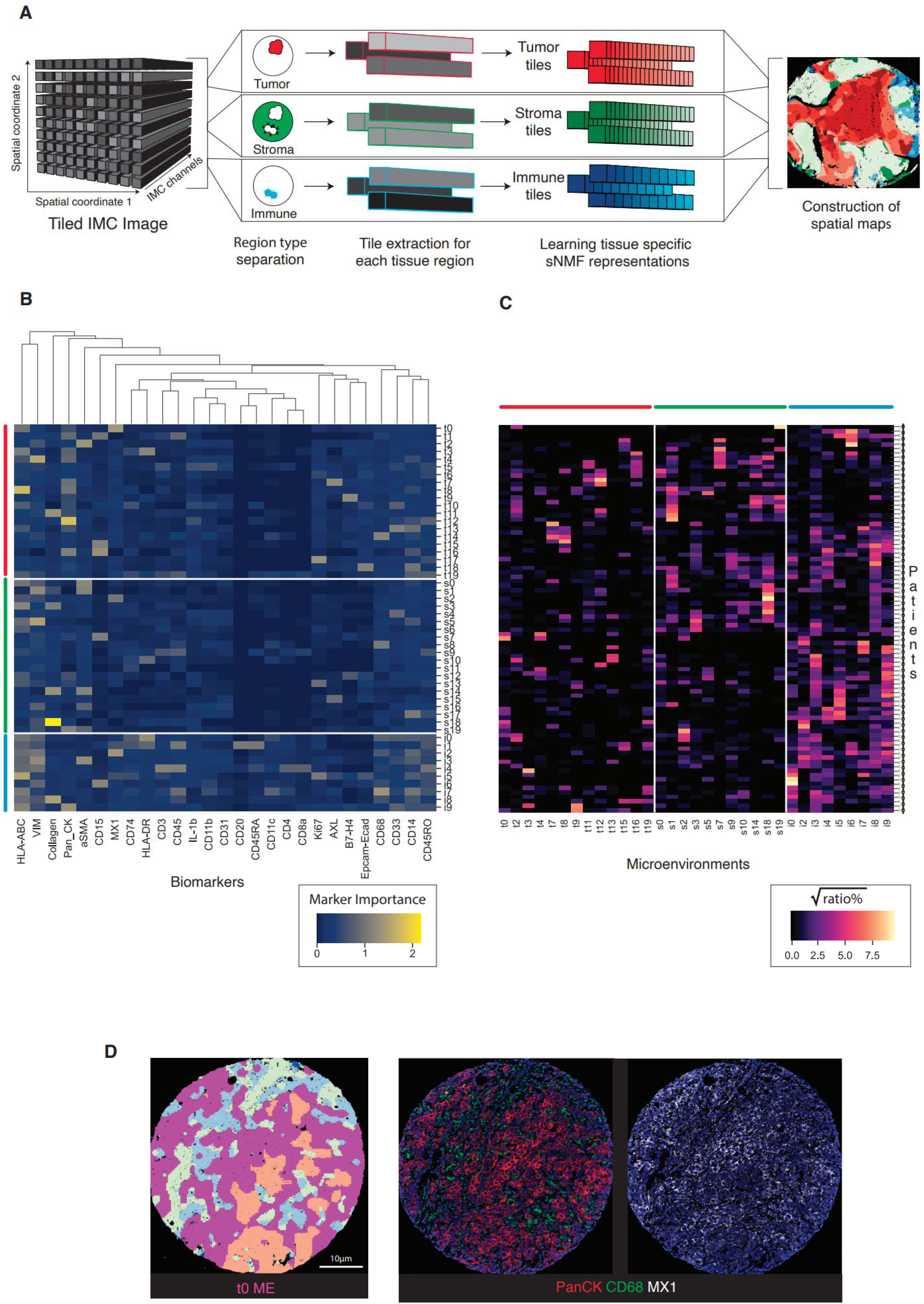
(A) overview of SparTile’s computational pipeline for ME identification: Three region types (tumor, stroma, and immune) are identified. Tiles within each region are encoded using sparse NMF. sparse NMF representations are brought back to pixel-level representations (methods), and each pixel is identified by the largest sparse NMF coefficient. (B) Importance of IMC markers for each SparTile cluster enables interpreting clusters as MEs. (C) Ratio of each ME across patients shows inter- and intra-patient heterogeneity. (B,C) The red, green, and blue bars denote tumor, stroma, and immune MEs, respectively. (D) Spatial mapping of SparTile clusters enables investigation of raw IMC images within each cluster. Magenta denotes the t0 ME and other colors denote other MEs within the TMA spot.

We found that SparTile clusters are interpretable and correspond to distinct MEs. The set of top IMC markers and marker pairs (Supplementary File 1), as well as marker importance scores (Fig. 2B) were used to interpret the ME corresponding to each cluster (Figs. 2D and 3A-C). For example, tumor cluster 0 (t0) is an MX1+ tumor-myeloid cell ME (Fig. 2D). PanCK, MX1, and myeloid markers, such as CD14 and CD68, are frequent among the top marker and marker pairs of t0, and are observed in raw IMC images in regions labeled as t0 (Fig. 2D). Furthermore, PanCK and MX1 have high importance in t0 (Fig. 2A).

### Individual tumor microenvironments correlate with survival

We observed that several MEs and expression patterns correlate with patient survival. t0 is an MX1+ tumor-myeloid cell ME. Remarkably, we found that the fraction of the spot classified as t0 correlates with worse survival (log-rank test p-value=0.04), even though the TMA spots are only 0.6 to 0.8mm-diameter tissue samples. t0 fraction was also predictive of survival risk in a Cox model accounting for clinical variables (methods, Fig. 3A, HR=5.25, CI=[1.32-9.18], p-value=0.01). This finding is consistent with previous studies showing that presence of myeloid cells and higher expression of MX1 in tumor ME is indicative of higher survival risk (Bianchini et al, 2022 and Aljohani et al, 2020). While MX1 in the tumor ME was associated with poor survival, the fraction of spot classified as i2, which is an MX1+ myeloid immune ME (Fig. 2B, Supplementary File 1), was not predictive of survival (log-rank p-value=0.66). Interestingly, the difference in t0 and i2 fraction (Supplementary Fig. 1, log-rank p-value=0.004) was a stronger predictor of survival than t0.

**Figure 3:**
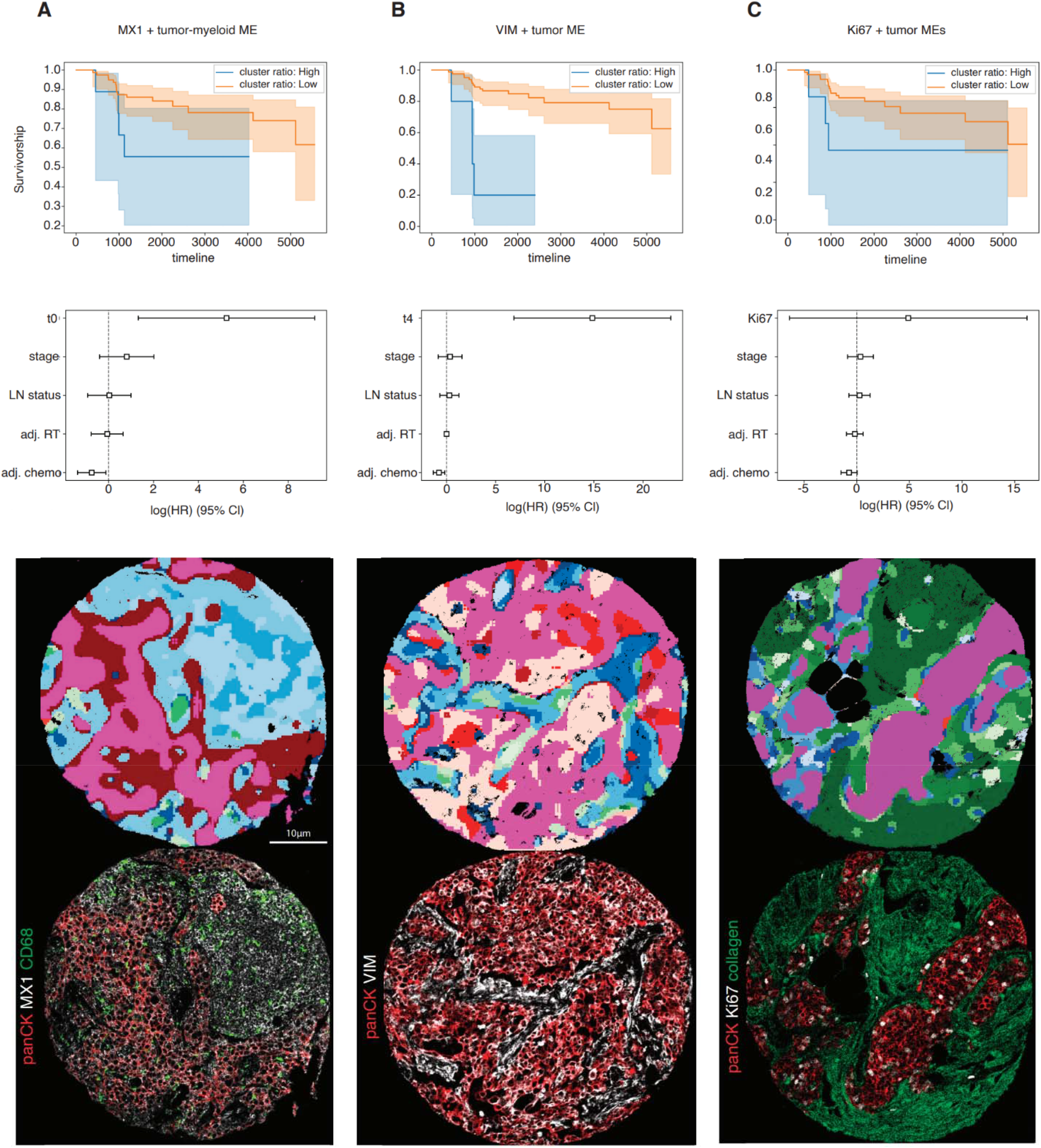
tumor MEs are predictive of survival risk: (top) Kaplan-Mier plots of patients stratified by ratios of distinct MEs. (middle) Hazard ratios of multivariate Cox models accounting for several clinical variables: stage, lymph node status, adjuvant chemotherapy (adj. chemo), and adjuvant radiotherapy (adj. RT). (bottom) Example regions of each ME. Magenta denotes the t0 ME and other colors denote other MEs within the TMA spot. (A) high ratio of MX1+ tumor-myeloid cell ME (t0) is predictive of poor survival. (B) High ratio of VIM+ tumors is predictive of poor survival. (C) High ratio of Ki67+ tumors individually correlates with poor survival, but it’s not statistically significant in a multivariate Cox model.

SparTile also revealed survival associations for t4, a vimentin+ tumor ME. Vimentin and panCK have high importance in t4 (Fig. 2B), and are prevalent in the list of top markers and top marker pairs (Supplementary File 1). We also frequently observed vimentin in manual inspection of t4 regions (Fig. 3B). Prior studies have shown that epithelial to mesenchymal transition is characterized by high expression of vimentin on tumor cells, suggesting t4 can potentially identify tumors with epithelial to mesenchymal transition (Grasset et al, 2020). Higher ratios of t4 correlates with poor survival (Fig. 3B, log-rank p-value=1.1e-5). Ratio of t4 was predictive of survival risk in a Cox model accounting for clinical variables (methods, Fig. 3B, HR=14.84, CI=[6.86-22.83], p-value<0.005).

Ki67 is frequently used as a marker of proliferation in tumor cells, and is an indicator of poor outcome (Keam et al, 2011). Importance scores (Fig. 2B), top IMC markers (Supplementary File 1), raw IMC images (Fig. 3C), and comparison of SparTile representations with different hyperparameters (Supplementary Fig. 2) show that SparTile clusters t6, t17, and s13 are contributors to the Ki67+ MEs. Higher fractions of Ki67+ MEs correlate with poor survival (Fig. 3C, log-rank p-value=0.052). Although Ki67+ status is an established marker of poor survival, we observed that Ki67+ ME fraction was not statistically significant in a multivariate Cox model accounting for clinical variables (methods, Fig. 3C, p-value=0.4).

Recent studies have emphasized the role of AXL in resistance to chemotherapy through promotion of proliferation and migration of cancer cells (Wium et al, 2021). t7 and s15 are the AXL+ MEs, and higher AXL+ ME fraction correlates with poor survival (log-rank p-value=0.03, Supplementary Fig. 2). The AXL+ ME fraction was not statistically significant in a multivariate Cox model accounting for clinical variables (methods, Supplementary Fig. 3, p-value=1).

While we observed that several tumor MEs correlate with survival, from a data-driven perspective, only t4 was statistically significant after correcting for multiple testing (methods). We believe this is due to the heterogeneity of TNBCs that result in a large number of distinct MEs with complex interactions.

### Individual immune microenvironments correlate with survival

A higher prevalence of lymphocytes has generally been considered to be positive for survival (Bianchini et al, 2022). Consistently, we observed that several lymphocyte-associated MEs within the stromal and immune region types correlate with longer survival. s9 is composed of CD8+ and CD45RO+ T-cells in collagen+ and aSMA+ stromal regions, and it is associated with longer survival (methods, T=0.03, log-rank p-value=0.07, Mann-Whitney U-test p-value=0.003, Supplementary Fig. 3). s9 fraction was a weak predictor of longer survival in a multivariate Cox model accounting for clinical variables (methods, Supplementary Fig. 3, HR=-21.98, CI=[-53.98, 9.94], p-value=0.18). i4 is a T-cell immune ME composed of both CD4+ and CD8+ T-cells (Supplementary File 1, Fig 2B). Higher i4 fraction correlates with longer survival (Supplementary Fig. 3, T=0.03, log-rank p-value=0.01, Mann-Whitney U-test p-value=2.1e-5). i4 fraction was statistically significant in a multivariate Cox model accounting for clinical variables (methods, Supplementary Fig. 3, HR=-31.62, CI=[-61.05, -2.19], p-value=0.04).

In contrast to the T-cell microenvironments, we observed a smaller effect size for B-cells (i1, Fig. 2B). i1 fraction weakly correlates with longer survival (methods, Mann-Whitney U-test p-value=0.03, log-rank p-value=0.26, Supplementary Fig. 3). We observed that B-cell MEs usually comprise a small fraction of the sample area (Supplementary Fig. 3). This exacerbates difficulties in representative quantification of B-cell ME prevalence from the small TMA spots, which may contribute to the weaker associations.

### Spatial distribution of tumors and myeloid cell markers is prognostic of survival

Because we observed distinct associations of myeloid protein prevalence with survival in tumor (t0 ME) and immune (i2 ME) regions, we hypothesized that *relative* spatial distributions between myeloid and tumor cells (Fig. 4A) might contain information to further improve survival predictions. To test this, we computed the total variation norm of the distribution of tumor and myeloid markers in each TMA spot, i.e. quantifying how separated tumor and myeloid cells are (methods). We observed a negative correlation (methods, Pearson correlation coefficient=-0.25) between large distance and large difference between ratios of t0 and i2 (methods, Fisher exact test p-value=0.036). Lower distance between tumors and myeloid markers are associated with poor survival (methods, Fig. 4B, p-value<0.01). Interestingly, we observed a higher expression of MX1 in images with low tumor-myeloid cell distance (Supplementary Fig. 4, Mann-Whitney U-test p-value=0.02). While previous studies have shown that intratumoral presence of myeloid cells, such as macrophages and neutrophils, correlates with poor outcome (Zheng et al, 2023; Qui et al, 2022), our results further quantify how myeloid infiltration affects survival. Spatial distance was statistically significant (p-value=0.02) in a multivariate Cox model accounting for several clinical variables (Fig. 4B). While stage has negative risk in this Cox model, it incurs no risk in a multivariate model without lymph node status (Supplementary Fig. 4,HR=-0.07, CI=[-0.73, 0.48], p-value=0.84), and has a weak positive risk in a univariate Cox model (HR=0.33, CI=[-0.28, 0.75], p-value=0.29). These results suggest that multicollinearity between variables may impact their hazard ratio in Cox models and emphasize the robustness of our spatial distance in predicting risk.

**Figure 4:**
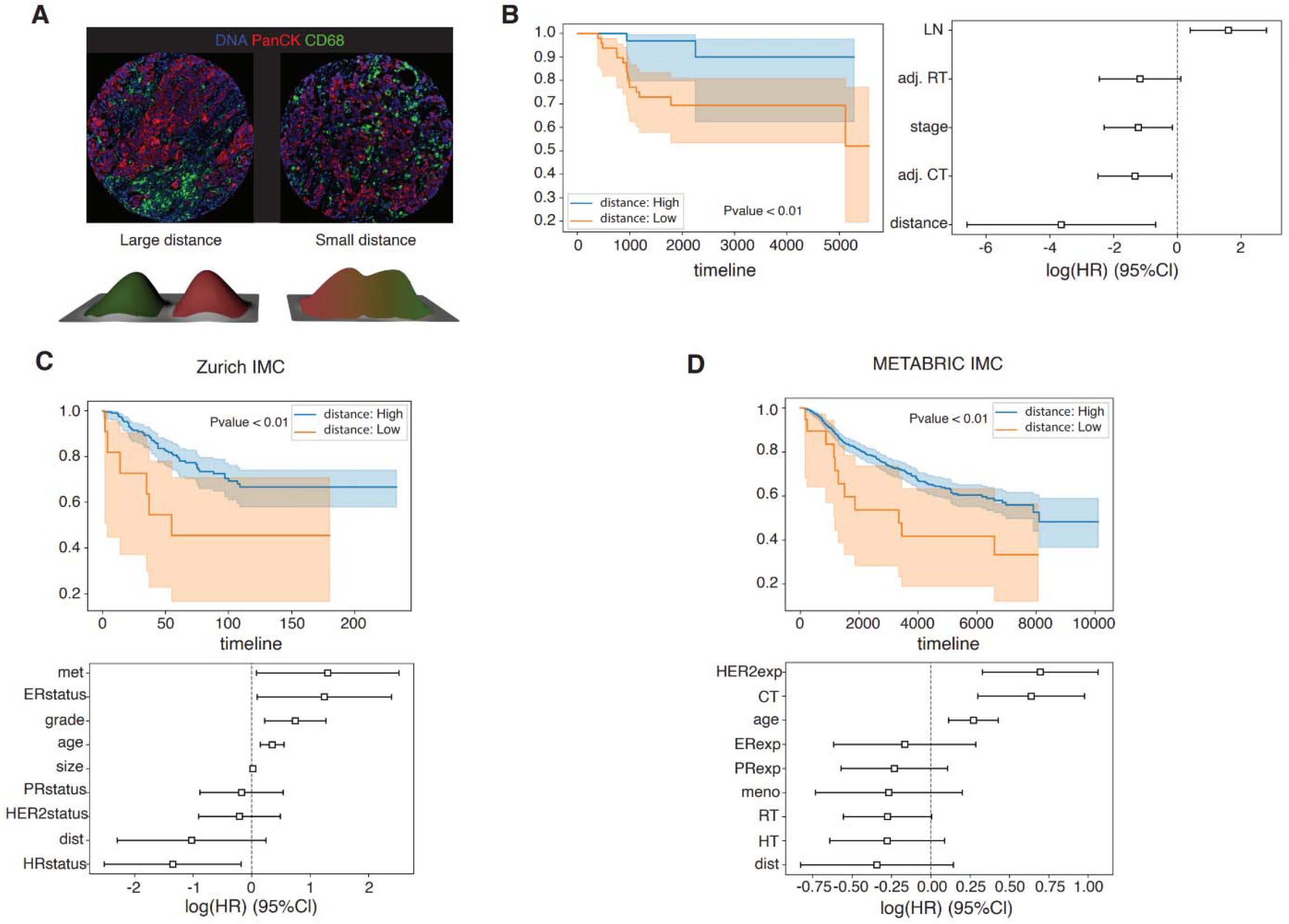
The spatial distribution of myeloid and tumor cells is predictive of survival. (a,top) Examples of IMC images with large (left) and small (right) distance between tumor and myeloid cells. (a,bottom) Example large and small distance tumor and myeloid cell marker distributions. (b) (left) large distance between tumor and myeloid cells is predictive of longer survival. (right) this association is statistically significant in a multivariate Cox model. (c,d) External validation of spatial distance as a predictor of survival on two external datasets: (c) Zurich IMC and (d) METABRIC IMC. (c) (left) The spatial distance of tumor and myeloid cells separates survival curves in the Zurich IMC data. (right) The spatial distance is an independent marker of survival in a multivariate Cox model accounting for the metastatic status (met), ER status, grade, age, tumor size, PR status, HER2 status, and HR status. (d) (left) The spatial distance of tumor and myeloid cells separates survival curves in the METABRIC IMC data. (right) The spatial distance is an independent marker of survival in a multivariate Cox model accounting for HER2, ER, and HR expression, age, treatment received (chemotherapy [CT], radio therapy [RT], and hormone therapy [HT]), and menopausal status (meno).

We used two external IMC datasets, the Zurich IMC dataset (Jackson et al, 2020) and the METABRIC IMC dataset (Ali et al, 2020), to further validate if the spatial distance between myeloid cells and tumors is indicative of survival in breast cancers (methods). Smaller distance correlates with poor survival in both datasets (Figs. 4c-d, Zurich IMC p-value<0.01, METABRIC IMC p-value<0.01). As both datasets are composed of multiple subtypes, we used a multivariate Cox model to account for subtype differences and other clinical variables. SparTile’s spatial distance was a weak independent prognostic marker of survival (Zurich IMC p-value=0.11, METABRIC IMC p-value=0.17) in both datasets. These results are interesting as both Cox models account for a wide range of clinical variables.

## Discussion

The tumor microenvironment is a complex and dynamic network of interactions involving cells, structural components, and signaling proteins. These spatial interactions are critical for understanding tumor evolution, therapeutic responses, and survival risks. SparTile offers a novel framework for studying these interactions by leveraging the spatial distribution of membrane, structural, and signaling proteins to distinguish diverse MEs. For example, t0 is a ME characterized by tumor and myeloid membrane markers and MX1 signaling protein.

While prior studies have demonstrated that marker prevalence can predict survival independent of spatial relations (Carter et al, 2023), SparTile advances this field by incorporating spatial structure at the tile level. This approach enables detailed analyses of multi-celltype driven survival associations. For example, SparTile summarized the spatial distribution of myeloid and tumor cell markers into a single metric, which effectively separated survival curves and demonstrated prognostic value in multivariate Cox models across three breast cancer datasets.

Segmentation-based methods have had encouraging results but face challenges, including error-prone cell segmentation and typing, as well as the computational burden of identifying and representing cellular neighborhoods. While graph neural networks have emerged as potential tools (Siao et al, 2024) their interpretability is limited, and incorporating structural and signaling proteins into their node representations remains complex. Additionally, the influence of network architecture and hyperparameter choices on the resulting representations is not fully understood.

SparTile addresses these limitations by employing a tile-based approach that encodes spatial relationships and facilitates the direct identification of distinct microenvironments. Compared to single-pixel analyses (eg. Liu et al, 2023), tiles provide richer contextual information, reducing the need for extensive preprocessing and hyperparameter tuning. SparTile has conceptual advantages over cell-cell contact-based metrics (e.g. Wang et al 2023), which fail to account for dynamic cellular states, such as the persistence of T-cell activation post-interaction with cancer cells. Because SparTile integrates such dynamics, it provides a more complete characterization of spatial information.

In summary, SparTile’s ability to incorporate membrane, structural, and signaling proteins into a cohesive analytical framework enhances the biological interpretability of multiplex imaging data. This capability enables more precise and comprehensive analyses, advancing our understanding of tumor biology and informing prognostic assessments.

## Methods

### Data collection

Tissue Microarrays (TMAs) were obtained from Yale School of Medicine (IRB#2019-088). TMA spots were 600 to 800 microns in diameter. Two adjacent 5-micron cuts were obtained for each spot. IMC was performed on TMA spots. Images corresponding to cell lines, relapsing or metastatic patients, or patients who received neoadjuvant therapy were removed from downstream analysis. 5 patients had two microarray spots, while all other patients had 1 spot. The images corresponding to 88 primary TNBC patients were used for downstream analysis. These patients were monitored, received adjuvant chemotherapy, adjuvant radiotherapy, or both adjuvant chemotherapy and radiotherapy.

Additionally, two publicly available breast cancer IMC datasets were used for analysis: METABRIC (Ali et al, 2020) and Zurich (Jackson et al, 2020). These datasets are composed of IMC images of TMAs of several breast cancer subtypes. Preprocessed IMC images of these datasets with no further preprocessing were used for downstream analysis.

### IMC data acquisition

TMA slides were incubated for 15 minutes at 58°C in a dry oven, deparaffinized in fresh histoclear, and rehydrated through a series of graded alcohols. Antigen retrieval was performed in a decloaking chamber (BioSB TintoRetriever) for 15 minutes at 95°C in citrate buffer, pH 6.0. After blocking in SuperBlock buffer (ThermoFisher Scientific), slides were incubated overnight at 4°C with a cocktail of metal-conjugated IMC-validated primary antibodies and described in Supplementary File 2. The following day, slides were washed twice in DPBS and counterstained with iridium intercalator (0.25 μmol/L) for 5 minutes at room temperature to visualize the DNA. After a final wash in Milli-Q water, the slides were air-dried for 20 minutes. The slides were then loaded on the Fluidigm Hyperion imaging mass cytometer. Each TMA spot was selected as a separate regions of interest using Fluidigm CyTOF Software (7.0) and ablated by the Hyperion. The resulting images were exported as 16-bit “.tiff” files using the Fluidigm MCDViewer software.

### Data preprocessing

The IMC images were preprocessed and denoised. Each IMC image was log normalized at the pixel level using log(1+x) transform. For each channel, pixels with values less than a threshold (T1) and connected components (8-connectivity) with size less than T2 were removed. The thresholds used for each channel are provided in Supplementary File 2.

### Visualization of IMC images

MCD Viewer was used to visualize raw IMC images from mcd files. For each IMC image and selected markers, the visualization parameters of MCD Viewer were selected to provide a reliable representation of marker distribution. These parameters are part of the MCD viewer software and were not used in the computational analysis via SparTile.

### Virtual H&E generation of IMC markers

DNA1, DNA3, CD3, CD20, and Ki76 were added to construct the nucleic channel, and aSMA, HLA-ABC, CD31, Collagen, PD1 and PDL-1 channels were added to construct the cytoplasmic channels. The nucleic and cytoplasmic channels were multiplied by a gain coefficient, and were fed to the falsecolor package (Serafin et al, 2020) to generate virtual H&E images. A 2D grid search was used to optimize gains of nucleic and cytoplasmic channels by minimizing L1 distance between median RGB values of virtual H&E images and a template image^*^ (Supplementary Fig. 5).

### Registering two IMC spots

We used the similarity transform of the Airlab package (v0.2.1, Sandkühler et al, 2018) to register grayscale virtual H&E images to each other. The IMC image of the second panel was transformed to match the coordinates of the first image. Max projection was used for IMC markers common between the two panels.

### Region type separation

IMC images were tiled using 20-micron overlapping tiles (75% overlap). Pixels with total IMC marker intensities of less than 1 were labeled as non-tissue, and tiles with less than 50% tissue pixels were removed. The average expression of IMC markers on each tile were used to compute the probability of each tile belonging to a tissue region type (tumor, stroma, immune). The following process, similar to Wang et al, 2023, was used to compute the probability of each tile belonging to each class. First, each region type was associated with its indicative markers. For example, tumor was associated with panCK and Epcam-Ecadherin markers. The full list of markers indicative of each region type is provided in Supplementary File 2. A quadratic link was used to compute the probability of each region type, i.e., for tile *T*, we have

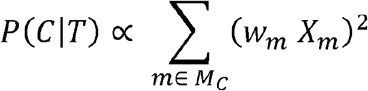

where *P*(*C*|*T*) is the probability tile *T* belongs to class *C* ∈ {*tumor, stroma, immune*}, *M*_*c*_ is the set of markers associated with class *C, w*_*m*_ is the weight for marker *m*, and *x*_*m*_ is the average expression of marker m in tile *T*. Weights are optimized by an iterative process. At each iteration, tiles were classified, and the gradient of the weight vector was used to update the weights. After training, given tile-level probabilities, the class probability vector of each pixel was calculated by averaging the class probabilities of all tiles containing the pixel.

### Tile representation learning

Given a set of tiles of a given size, the pixel-level tissue region type probability maps were used to obtain tile-level probabilities by averaging pixel-level probabilities over each tile. Then, each tile was associated with the class with the largest probability. For tiles belonging to each region type, average expression of IMC markers and IMC marker pair colocalizations were calculated. To compute colocalization of two IMC markers, first each marker was dilated by disk (K=5). Then, dilated images of the two IMC markers were multiplied at the pixel level, and the average value of the product channel over the tile was saved. Given the average expression of IMC markers and marker pairs, sparse NMF (sNMF) was used to reduce dimensionality of encoded tiles to D=10,15, or 20 dimensions.

IMC images were tiled using overlapping tiles with the following sizes: 20, 30, and 40 microns. The sNMF analysis described above was performed for each tile size. For each tissue region type, 9 sNMF models corresponding to the 3 tile sizes (20, 30, and 40) and 3 dimensionality reduction parameters (D=10,15,20), were computed.

### Clustering tiles in each region type

First, tissue pixels, i.e., those with total IMC intensity larger than 1, were classified as a tumor, stroma, or immune. Given tile-level representations, pixel-level representations were obtained by averaging over the tiles containing the pixel. For each pixel, the sNMF model corresponding to the pixel region type was used for all tiles containing it. Finally, each pixel was associated with the sNMF component with the largest coefficient.

### Importance scores of each cluster

An importance score was computed for IMC markers of each sNMF cluster:

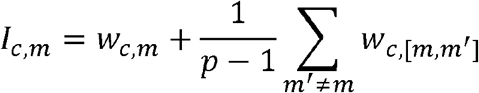

where *I*_*c*,*m*_ is the importance of marker *m* in sNMF cluster *c, w*_*c*,*m*_ is the weight of marker *m* in sNMF component *c*,*w*_*c*,, *[m*,*m’]*_ is the weight of marker pair *m* and *m’* in sNMF component *c*, and *P* is the total number of IMC markers used for analysis.

### Interpretation of SparTile clusters as MEs

Top IMC markers and marker pairs of an sNMF cluster, importance scores of IMC markers, cross-referencing clusters across representations of different sNMF models, and confirmation of marker presence in raw IMC data of a cluster were used to interpret sNMF clusters and identify the MEs they encode. These interpretations were additionally used to select the hyperparameters of SparTile. We used tile sizes of 30 microns (D=20), 30 microns (D=20), and 20 microns (D=10) for tumor, stroma, and immune regions, respectively.

### Association of SparTile clusters with survival

The fraction of tissue pixels corresponding to each SparTile cluster was computed for each patient. Unless otherwise stated, a ratio threshold of 0.05 was used to separate patients into high and low cluster ratios. Lower thresholds were used for less prevalent SparTile clusters. The log rank test was used to compare Kaplan-Meier (KM) plots of patients in high and low ratio groups. Additionally, patients with overall survival less/more than 3 years were labeled as short/long survival. Mann-Whitney-U test was used to compare ratio of sNMF clusters between the two groups as an additional test.

### Spatial distance of tumors and myeloid cell markers

PanCK was used as the tumor marker, and the sum of CD68, CD11b, CD14, and CD33 markers was used as the myeloid marker. First, patients with low average expression of tumor or myeloid markers were removed (T=0.05). The remaining 80 patients were used in downstream analysis. The tumor and myeloid markers were blurred using a Gaussian Kernel with a bandwidth of 50 pixels. Afterwards, the blurred images were normalized to have a unit volume, representing the 2D distributions of tumor and myeloid markers. The total variation norm between these two distributions was used as the distance:

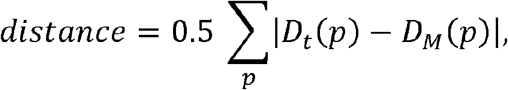

where *D*_*t*_ and *D*_*m*_ are the distributions of tumor and myeloid markers, and *P* denotes the pixels in an IMC image. A threshold of 0.25 was used to separate images with high/low distance.

For the METABRIC IMC images, we used panCK as the tumor marker, sum of CD68, CD163, and CD11c as myeloid markers, used a threshold of 0.1 for selecting images staining positive for tumor and myeloid cell markers, and a threshold of 0.15 to separate images with high/low distance between tumor and myeloid markers. For the Zurich IMC dataset we used sum of panCK, CK7, CK8/18, CK19, CK5, and KRT14 as the tumor marker, CD68 as myeloid marker, used a threshold of 0.2 for selecting images staining positive for tumor and myeloid cell markers, and a threshold of 0.15 to separate images with high/low distance between tumor and myeloid cell markers.

### Multivariate Cox proportional hazard model analysis

The lifelines python package (v0.27.4) (Davidson-Pilon, 2019) was used to train multivariate Cox models for a set of independent variables. The p-values and confidence intervals of the trained Cox model are reported. For Multivariate Cox models using ratios of SparTile MEs we used elastic-net regularization (penalty=0.01, l1_ratio=0.5) for stable parameter estimation. No regularization was used for other Cox models.

### Correlation between Spatial distance and ratio of MEs

The difference between t0 and i2 was used for analysis. Patients with low fraction of t0 or i2 were removed. Patients were labeled as distance high/low based on the spatial distance, and difference high/low based on difference of t0 and i2. Fisher exact test was used to test if there is a correlation between spatial distance and difference of t0 and i2.

### Computational Implementation

All computational analyses were performed in python. The following packages were used: imctools (v2.1.8) (Windhager et al, 2023), numpy (v1.26.0) (Harris et al, 2020), scipy (v1.8.1) (Virtanen et al, 2020), falsecolor (Serafin et al, 2020), Airlab (v0.2.1) (Sandkühler et al, 2018) scikit learn (v1.1.2) (Pedregosa et al, 2011), matplotlib (v3.5.2) (Huner, 2007), seaborn(v0.12.0) (Waskom, 2021), lifelines (v0.27.4) (Davidson-Pilon, 2019), and pandas (v1.4) (McKinney, 2010).

## Acknowledgements

Research reported in this publication was supported by NIH grant R01CA230031, NIH grant RO1CA219880, and NIH/NCI grant P30 CA034196.

## Data and code availability

The data and code associated with the manuscript will be made available upon publication.

## Supplementary Figures

**Supplementary Figure 1:**
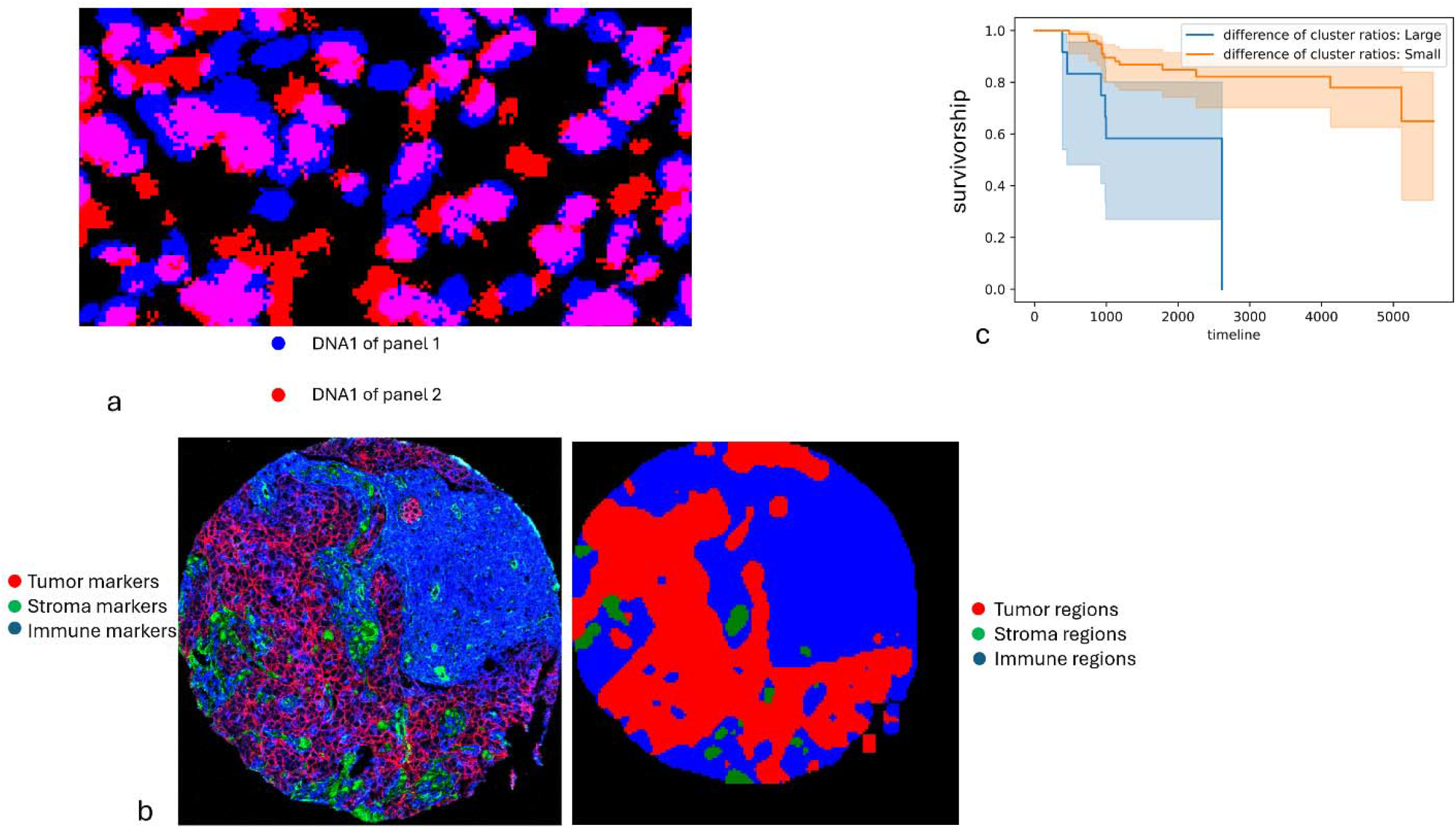
(a) Snapshot of DNA1 markers of two registered images. While most cells match, cells being present in only 1 image, difference of nucleus shape between the two images, and registration errors impede reliable cell segmentation. (b) Regions rich in tumor, stromal, and immune markers are separated. Left: expression of markers indicative of tumor cells (red, panCK and Epcam-Ecadh), stroma (green, Collagen, aSMA, and vimentin), and immune cells (blue, CD45, CD45RA, CD45RO, CD33, CD3, CD4, CD8a, CD20, CD11b, CD14, CD68, HLA-DR). Right: Reigns predicted to be dominated by tumor cells, stromal regions, and immune regions outside tumor. (c) The difference of ratios of t0 and i2 separates survival curves (T=0.01).

**Supplementary Figure 2:**
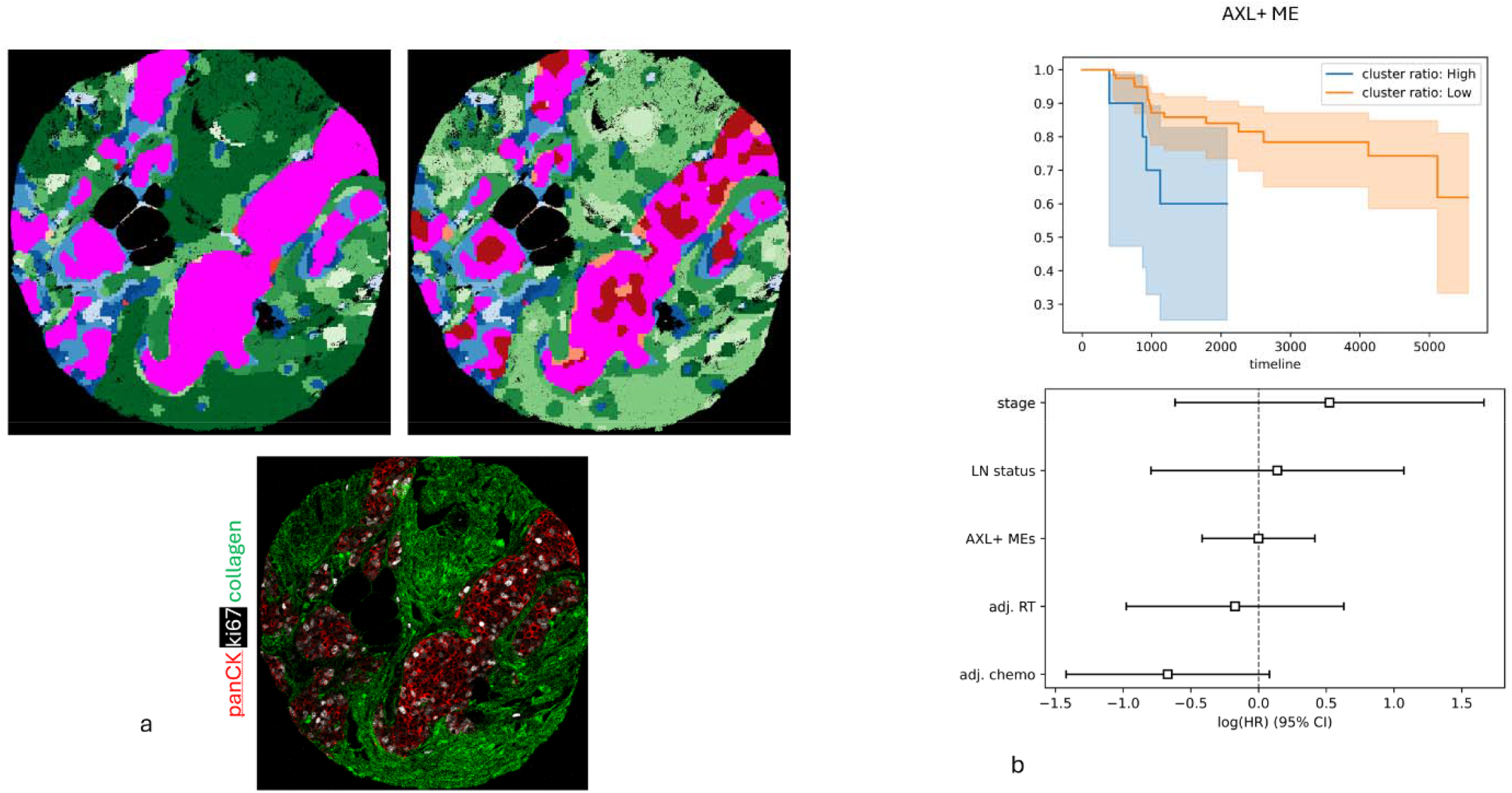
(a) Top left, t6 ME highlighted in magenta for T=30, top right, t6 ME highlighted in magenta for T=40, bottom, raw IMC image. (b) Higher ratio of AXL+ MEs correlates with shorter survival, but it is not an independent marker of risk in multivariate Cox models.

**Supplementary Figure 3:**
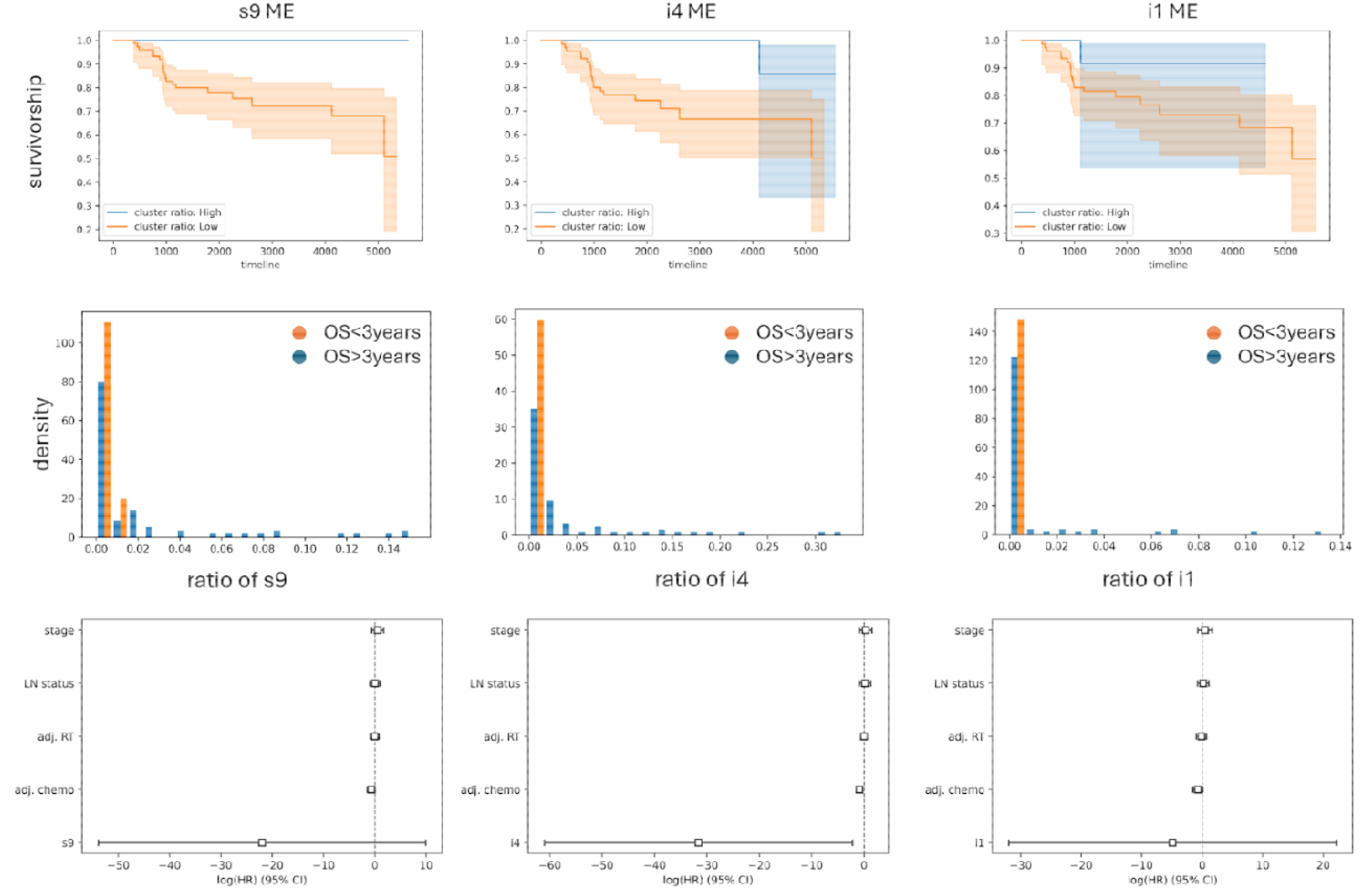
(top,left) higher ratio of s9 correlates with longer survival. (middle,left) A subset of patients with overall survival (OS) > 3 years have higher ratios of s9 (>0.02), while all patients with short survival (OS<3 years) have small ratios of s9 (<0.02). (bottom, left) The ratio of s9 has a negative hazard ratio, but not an independent marker of survival in a multivariate Cox model. (top,middle) higher ratio of i4 correlates with longer survival. (middle,middle) A subset of patients with OS> 3 years have higher ratios of i4, while all patients with short survival (OS<3 years) have small ratios of i4. (bottom, middle) The ratio of i4 has a negative hazard ratio, and is an independent marker of survival in a multivariate Cox model. (top,right) higher ratio of i1 correlates with longer survival. (middle,right) A subset of patients with OS> 3 years have higher ratios of i1, while all patients with short survival (OS<3 years) have small ratios of i1. (bottom, left) The ratio of i1 has a negative hazard ratio, but not an independent marker of survival in a multivariate Cox model.

**Supplementary Figure 4:**
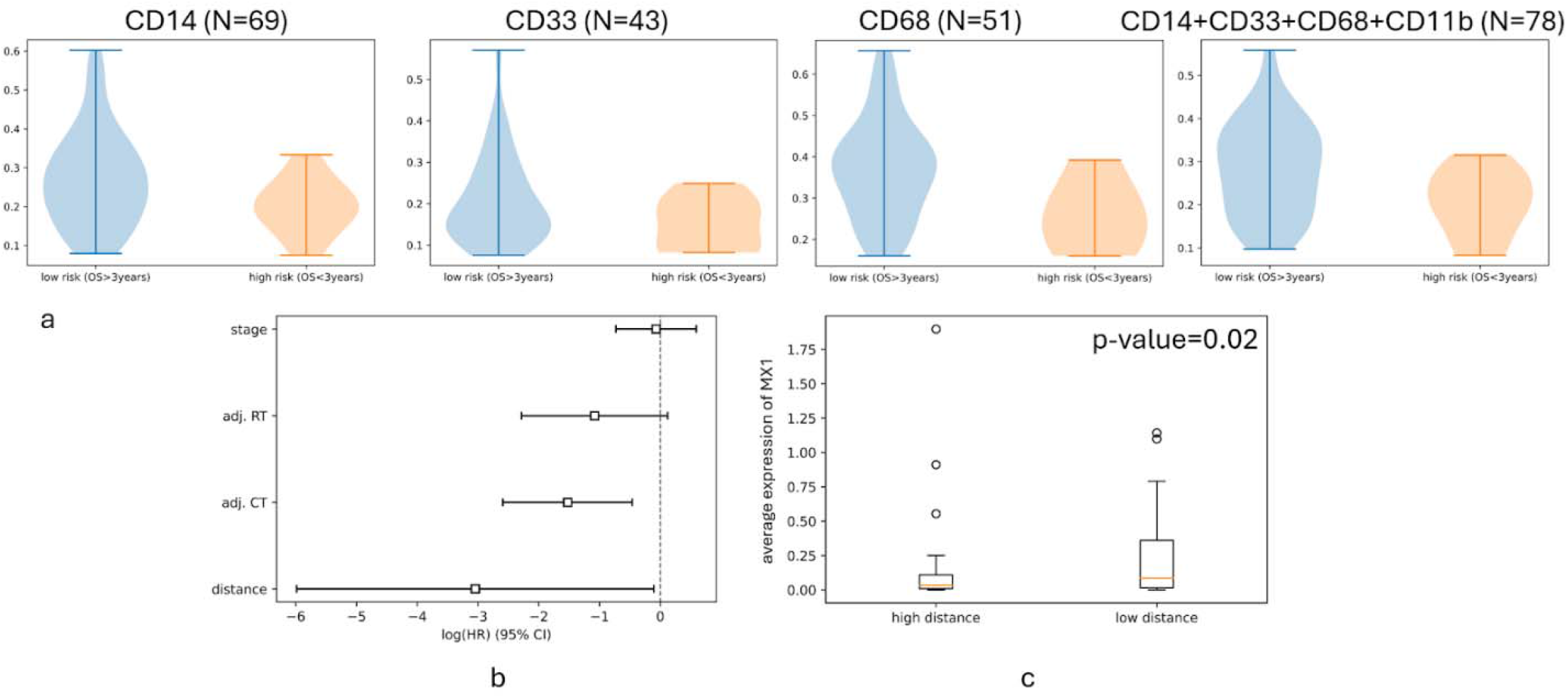
(a) The spatial distance of panCK and several myeloid cell markers. Consistently, large distance between panCK and myeloid cell markers is indicative of longer overall survival. (b) In a multivariate Cox model not accounting for the lymph node status, distance of tumor and myeloid cells is an independent predictor of survival risk, while stage is not an independent predictor of survival risk. (c) MX1 has a higher average expression in IMC images with low distance between tumor and myeloid cells.

**Supplementary Figure 5:**
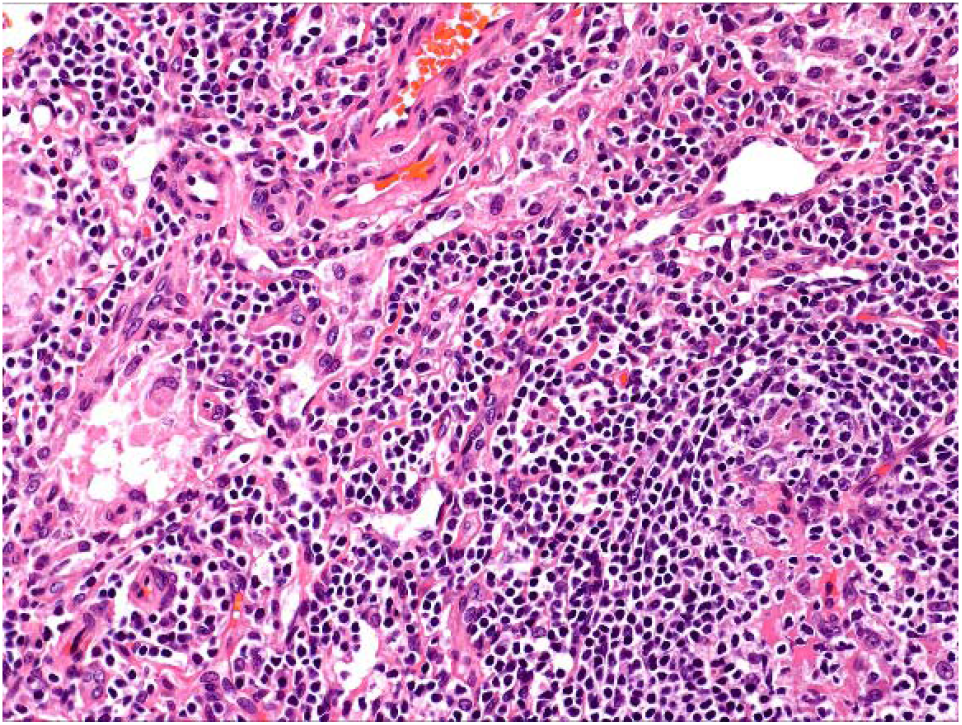
The template image used to generate virtualized H&E images (obtained from https://www.pathologyoutlines.com/topic/liveradenovirus.html).

## Footnotes

* Obtained from: https://www.pathologyoutlines.com/topic/liveradenovirus.html

## References

Ali, H.R., Jackson, H.W., Zanotelli, V.R.T. et al. Imaging mass cytometry and multiplatform genomics define the phenogenomic landscape of breast cancer. Nat Cancer 1, 163–175 (2020).

Aljohani, A.I., Joseph, C., Kurozumi, S. et al. Myxovirus resistance 1 (MX1) is an independent predictor of poor outcome in invasive breast cancer. Breast Cancer Res Treat 181, 541–551 (2020).

Bianchini, G., De Angelis, C., Licata, L. et al. Treatment landscape of triple-negative breast cancer — expanded options, evolving needs. Nat Rev Clin Oncol 19, 91–113 (2022).

Carter, J. M., Chumsri, S., Hinerfeld, D.A. et al. Distinct spatial immune microlandscapes are independently associated with outcomes in triple-negative breast cancer. Nat. Commun. 14, 2215 (2023)

Chen, Z., Fang, Z., Ma, J. Regulatory mechanisms and clinical significance of vimentin in breast cancer. Biomedicine & Pharmacotherapy. 1, 133:111068 (2021).

Davidson-Pilon, C. lifelines: survival analysis in Python. J. Open Source Software, 4:40, (2019).

Grasset, E.M., Dunworth, M., Sharma, G. et al. Triple-negative breast cancer metastasis involves complex epithelial-mesenchymal transition dynamics and requires vimentin. Science translational medicine, 14:656 (2022).

Harris, C.R., Millman, K.J., van der Walt, S.J. et al. Array programming with NumPy. Nature 585, 357–362 (2020).

Hunter, J. D. “Matplotlib: A 2D graphics environment.” Computing in science & engineering 9(03), 90–95 (2007).

Jackson, H.W., Fischer, J.R., Zanotelli, V.R.T. et al. The single-cell pathology landscape of breast cancer. Nature 578, 615–620 (2020).

Keam, B., Im, S. A., Lee, K. H. et al. Ki-67 can be used for further classification of triple negative breast cancer into two subtypes with different response and prognosis. Breast Cancer Res 13, R22 (2011).

Kuburich, N. A., den Hollander, P., Pietz, J.T., Mani, S. A. Vimentin and cytokeratin: Good alone, bad together. In: Seminars in cancer biology, 86, 816–826 (2022). Academic Press.

Liu, C.C., Greenwald, N.F., Kong, A. et al. Robust phenotyping of highly multiplexed tissue imaging data using pixel-level clustering. Nat Commun 14, 4618 (2023).

McKinney, W. “Data structures for statistical computing in Python.” SciPy 445.1, 51–56 (2010).

Pedregosa, F., Varoquaux, G., Gramfort, A. et al. Scikit-learn: Machine learning in Python. J. Machine Learning Research, 12, 2825–2830 (2011).

Qiu X., Zhao T., Luo, R., Qiu, R., Li, Z. Tumor-associated macrophages: key players in triplenegative breast cancer. Frontiers in Oncology. 14, 12:772615 (2022).

Sandkühler, R., Jud, C., Andermatt, S., Cattin, P. C. AirLab: autograd image registration laboratory. arXiv preprint 1806.09907. (2018).

Serafin, R., Xie, W., Glaser, A. K., Liu, J.T. FalseColor-Python: a rapid intensity-leveling and digital-staining package for fluorescence-based slide-free digital pathology. Plos one. 1,15(10):e0233198 (2020).

Shiao, S.L., Gouin, K.H., Ing, N., et al. Single-cell and spatial profiling identify three response trajectories to pembrolizumab and radiation therapy in triple negative breast cancer. Cancer cell, 42(1), 70–84 (2024).

Virtanen, P., Gommers, R., Oliphant, T.E. et al. SciPy 1.0: fundamental algorithms for scientific computing in Python. Nat Methods 17, 261–272 (2020).

Vora, H. H., Patel, N. A., Rajvik, K. N., et al. Cytokeratin and Vimentin Expression in Breast Cancer. Int. J. Biol Markers. 24(1), 38–46, (2009).

Wang, V.G., Liu, Z., Martinek, J. et al. Computational immune synapse analysis reveals T-cell interactions in distinct tumor microenvironments. Commun Biol 7, 1201 (2024).

Waskom, M. L. seaborn: statistical data visualization. J. Open Source Software, 6(60), 3021 (2021).

Windhager, J., Zanotelli, V.R.T., Schulz, D. et al. An end-to-end workflow for multiplexed image processing and analysis. Nat Protoc 18, 3565–3613 (2023).

Wium, M., Ajayi-Smith, A. F., Paccez J. D., Zerbini, L. F. The Role of the Receptor Tyrosine Kinase Axl in Carcinogenesis and Development of Therapeutic Resistance: An Overview of Molecular Mechanisms and Future Applications. Cancers (Basel). 25,13(7):1521 (2021)

Zagami, P., Carey, L. A. Triple negative breast cancer: Pitfalls and progress. npj Breast Cancer 8, 95 (2022).

Zheng, C., Xu, X., Wu, M. et al. Neutrophils in triple-negative breast cancer: an underestimated player with increasingly recognized importance. Breast Cancer Res 25, 88 (2023).

